# A Novel Angiogenesis Role of GLP-1(32-36) to Rescue Diabetic Ischemic Lower Limbs via GLP-1R-Dependent Glycolysis

**DOI:** 10.1101/2023.06.01.543344

**Authors:** Yikai Zhang, Shengyao Wang, Qiao Zhou, Yepeng Hu, Yi Xie, Weihuan Fang, Changxin Yang, Zhe Wang, Shu Ye, Xinyi Wang, Chao Zheng

## Abstract

Glucagon-like peptide 1 (GLP-1) improves angiogenesis, but the mechanism remains unclear. To address this question, we conducted a metabolomics analysis in bone marrow-derived endothelial progenitor cells (EPCs) isolated from T1DM mice treated with or without GLP-1(32-36) amide, an end-product of GLP-1. GLP-1(32-36) treatment recovered glycolysis. In addition, GLP-1(32-36) treatment rescued diabetic ischemic lower limbs and EPCs dysfunction by regulating PFKFB3-driven glycolytic flux capacity and mitochondrial dynamics. The effects of GLP-1(32-36) were abolished in mice lacking a functional GLP-1 receptor (*Glp1r*^-/-^), which could be partially rescued in EPCs transiently expressing GLP-1R. GLP-1(32-36) treatment activated the eNOS/cGMP/PKG pathway, increased glycolysis, and enhanced EPCs angiogenesis. Taken together, these findings suggest that GLP-1(32-36) could be used as a monotherapy or add-on therapy with existing treatments for DPAD.

**Graphical abstract:** 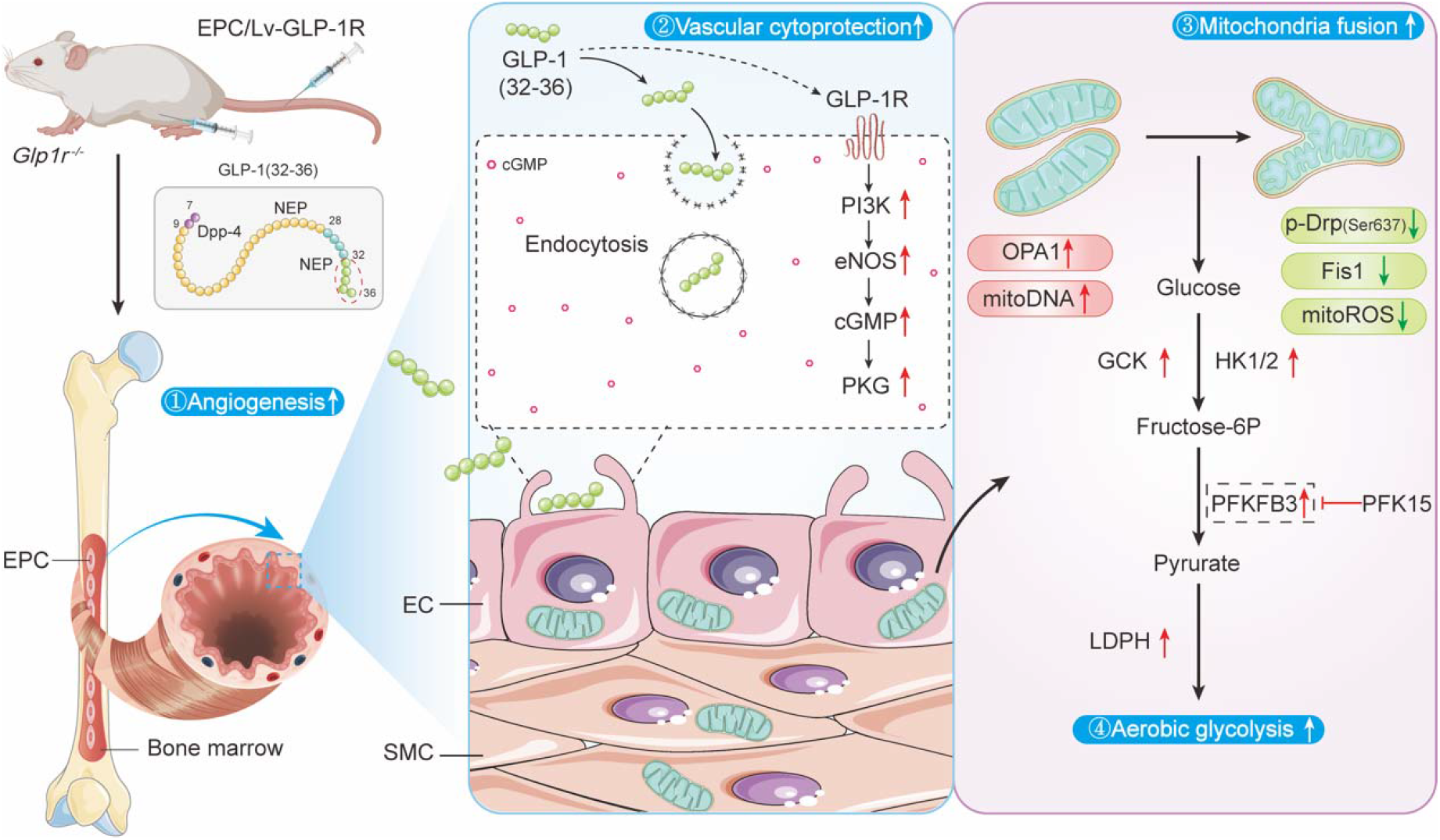

## 1. Introduction

Peripheral arterial disease (PAD), a severe chronic complication of diabetes, is characterized by the narrowing and occlusion of arteries supplying the lower extremities (1). Although PAD typically presents as claudication, it can progress to critical limb ischemia and may eventually require amputation (2). Given that diabetic patients are at a fourfold greater risk of developing PAD than the general population, there is likely a close relationship between hyperglycemia and vascular complications (3). Recent studies reveal that incretins such as glucagon-like peptide 1 (GLP-1) play a role in modulating angiogenesis beyond their function in glycemic control (4, 5).

GLP-1 is a naturally occurring hormone that plays a vital role in regulating glucose homeostasis by stimulating insulin secretion (6). It is produced by enteroendocrine L cells located in the distal ileum and colon, and is found in two molecular forms, GLP-1 (7-37) and GLP-1 (7-36) amide, which bind to a specific G-protein coupled receptor called GLP-1R (7, 8). However, they are rapidly degraded by the enzyme dipeptidyl peptidase-4 (DPP-4) after their release into the bloodstream, resulting in the formation of amino-terminally truncated peptides, such as GLP-1(9-37) and GLP-1(9-36) amide (9). GLP-1(9-36) amide can enter cells by penetrating the cell membrane, where it is internally cleaved in the C-terminal region by an intracellular endopeptidase such as neutral endopeptidase 24.11 (NEP 24.11), leading to the production of the nonapeptide GLP-1 (28-36) and the pentapeptide GLP-1(32-36) amide (10) (Fig. 1A).

**Figure 1:**
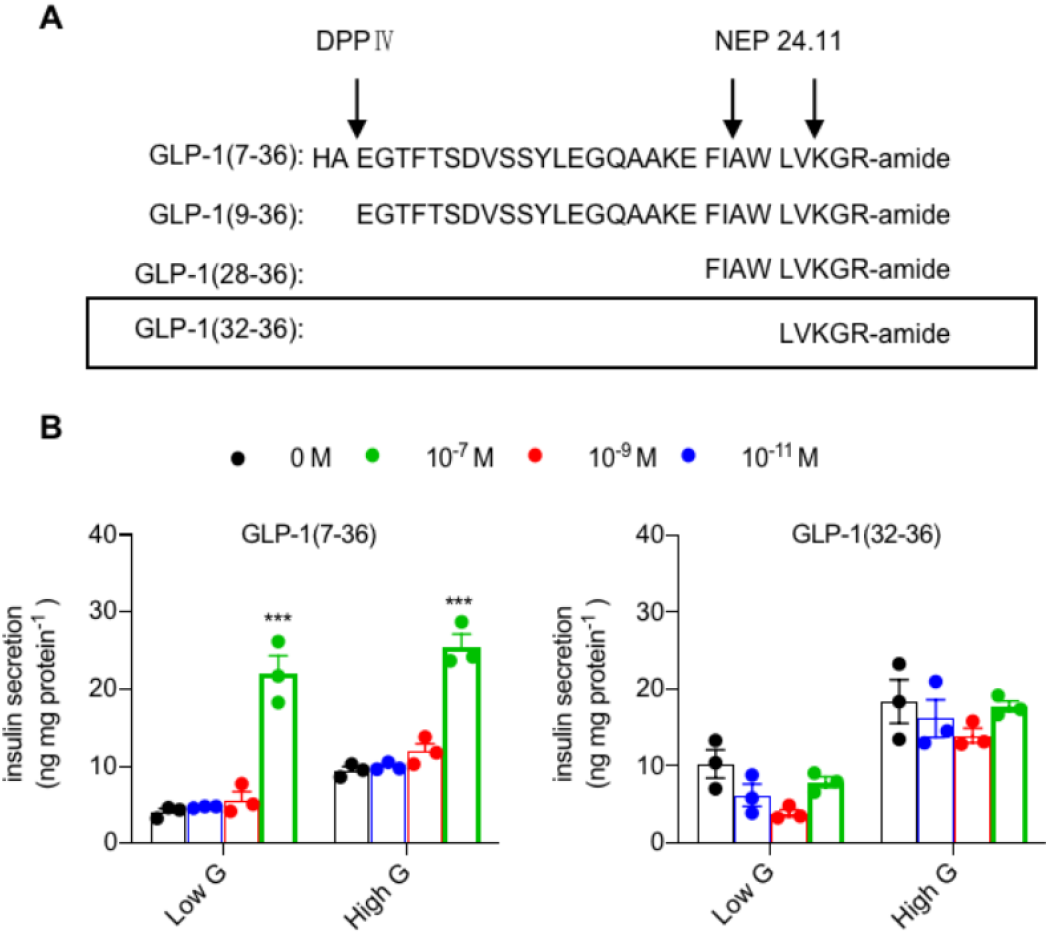
GLP-1 metabolites i11j1ue11t different dose glucose-stimulated i11s11/i11secretion of INS-I cells. (A)Formation of the C-terminal peptides GLP-1(7-36), GLP l (9-36), GLP-1(28-36), and GLP-1(32-36) amides. (B) INS-I cells were treated with either GLP-1(7-36) or GLP-1(32-36) with different dose (0 M, 10·^11^ M, 10· ^9^ M, 10·^7^ M), and 48 h later, insulin secretion was measured under low glucose (2 111M) or high glucose (33 111M) concentrations. Cells were equilibrated with KRBH buffer and stimulated with 2.8 111M, 20 111M glucose and with 40 111M KC/ (primary islet cells). Mean ±SEM from independent experiments (B; 11=3). Statistical significance was determined by I-way ANOVA with post hoc Tukey multiple comparisons test. *P<0.05; **P<0.01 and ***P<0.001.

Several GLP-1 peptides and their metabolites are reported to have a cardiovascular protective effect. For example, GLP-1(9-36) has been shown to improve human aortic endothelial cell viability in response to hypoxia via a NO-dependent mechanism (11). GLP-1(9-36) has also been found to reduce high glucose-induced mitochondrial ROS generation in human endothelial cells (12). GLP-1(28-36) activates the AC-cAMP signaling pathway, changes the metabolic status of vascular cells, and plays a role in cardiovascular protection (13). GLP-1 (32-36), the major end-product of GLP-1 proteolysis in addition to the nonapeptide, has been found to decrease body weight, increase energy consumption, and reduce β-cell apoptosis in obese mice (14, 15). However, whether GLP-1 (32-36) has a beneficial effect on diabetic vascular endothelial injury remains unclear.

By binding to its receptor (GLP-1R) expressed in various organs, including pancreatic islets, heart, lungs, and brain stem, GLP-1 activates the cAMP-dependent signal transduction pathway to promote glucose-dependent insulin secretion by β-cells and improve nutrient utilization in peripheral organs (16). While GLP-1 metabolites have been found to exert beneficial effects when administered parenterally, their mechanisms of action are not fully understood (17-20). GLP-1 has been shown to protect against heart failure through the GLP-1R-mediated eNOS/cGMP/PKG pathway rather than the cAMP/PKA pathway (21). The mode of GLP-1(32-36) entry and its possible signaling mechanisms, if ever exist, remain unknown.

It has been proposed that endothelial cells (ECs) play a critical role in maintaining vascular homeostasis and promoting angiogenesis by relying on glycolysis to produce more than 80% of their ATP (22-24). A recent study suggests that GLP-1 can regulate astrocytic glycolysis, which may contribute to its neuroprotective effects in Alzheimer’s disease (25). Therefore, it is reasonable to hypothesize that GLP-1(32-36) may enhance glycolytic flux in ECs, thereby altering vessel sprouting and promoting angiogenesis.

In this study, we demonstrate that GLP-1(32-36) administration has a direct effect on diabetic lower limb ischemia. We also demonstrate that GLP-1(32-36) has a causal role in improving fragile mitochondrial function and metabolism via the GLP-1R-mediated pathway, independent of its insulinotropic action. Specifically, we found that GLP-1(32-36) promotes angiogenesis in endothelial progenitor cells (EPCs) exposed to high glucose and enhances blood perfusion in ischemic tissues in STZ-induced type 1 diabetic mice with hindlimb ischemia (HLI). We also show that GLP-1(32-36) improves mitochondrial dynamics and rescues glycolysis mediated by 6-phosphofructo-2-kinase/fructose-2,6-bisphosphatase 3 (PFKFB3). We further demonstrate that GLP-1(32-36) rescues diabetic ischemic lower limbs by activating the GLP-1R-dependent eNOS/cGMP/PKG pathway. Our findings provide novel insights into the mechanisms underlying the beneficial effects of GLP-1(32-36) on DPAD and highlight its potential therapeutic value for non-diabetic patients due to its angiogenic effect that is independent of insulin regulation.

## 2. Results

### 2.1 GLP-1 (32-36) promotes blood perfusion and angiogenesis post-HLI in type 1 diabetic mice independent of insulinotropic actions

We found that GLP-1(32-36) treatment protected human umbilical vein endothelial cells (HUVECs) from high glucose (HG)-induced reduction in tuber formation (Supplementary Fig. 1), indicating improved HUVEC integrity and function. Unlike GLP-1(7-36), GLP-1(32-36) does not stimulate insulin secretion from insulinoma 1 (INS-1) cell under either high or low glucose conditions, suggesting that the angiogenic capability of this pentapeptide is independent of insulinotropic action (Fig. 1B). To investigate whether GLP-1(32-36) promotes angiogenesis in vivo, we used STZ-induced type 1 diabetic mice (T1DM) as a murine model of unilateral hind limb ischemia (HLI) to examine the therapeutic potential of GLP-1(32-36) on angiogenesis according to previous protocols (26-28) (Supplemental Fig. 2). The mice were treated with GLP-1(32-36) (1μmol/kg/d), GLP-1(7-36) (1μmol/kg/d), or PBS by daily intraperitoneal (i.p.) infusion and the blood flow recovery was evaluated by using a PeriCam Perfusion Speckle Imager (PSI) at day 0, 3, 7, 14, 21, 28 days after HLI surgery (Supplementary Table. 1). GLP-1(7-36) here was used as a control for angiogenic drug (28). GLP-1(32-36) treatment markedly recovered blood flow, companied with improved neovascularization in ischemic tissue (Fig. 2A). The effect of the pentapeptide in promoting neovascularization was characterized by enhanced CD31 expression in ischemic gastrocnemius muscle, as shown by Western blotting and immunofluorescence staining (Fig. 2B-E). To investigate the effect of GLP-1(32-36) on EPC mobilization in response to tissue ischemia, we examined the numbers of double positive Sca-1^+^/Flk-1^+^ cells in mononuclear fraction of peripheral blood from T1DM mice by flow cytometry. Administration of GLP-1(32-36) into tissue ischemia T1DM mice substantially augmented EPC mobilization on day 3 and peaked on day 7 after HLI (Fig. 2F-G). These results demonstrate that GLP-1(32-36) is superior to GLP-1(7-36) in rescuing angiogenic function and blood perfusion in ischemic limb of STZ-induced diabetic mice without an effect on insulin secretion.

**Figure 2:**
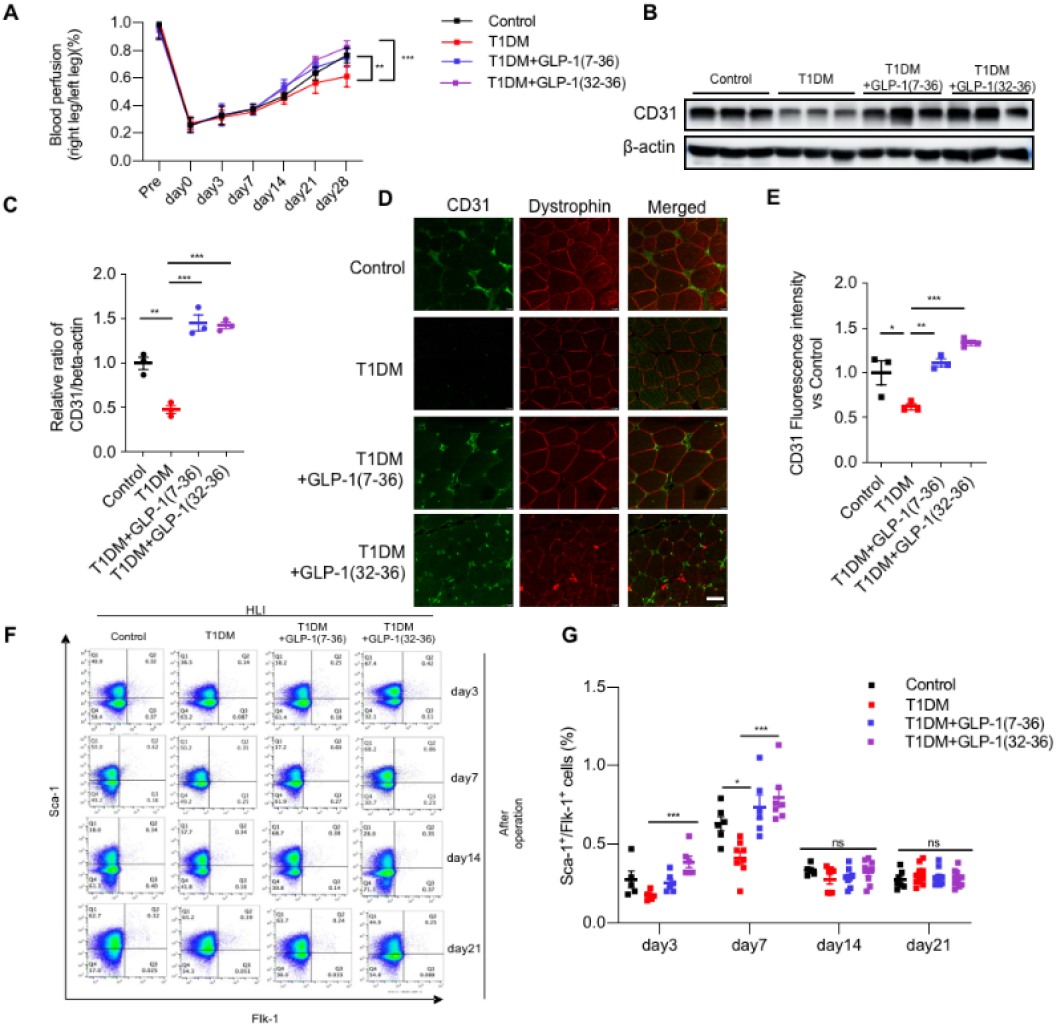
GLP-K32-36) improves blood perfusion, angiogenesis, and endothelial progenitor cell mobilization in ischemic hindlimbs of type 1 diabetic mice (T1DM). The proangiogenic effect of GLP-1 (32-36) in diabetic ischemic tissues was investigated in T1DM mice. GLP-l(7-36) is as a full-length peptide control. (A) blood flow reperfusion was assessed by Doppler laser ultrasound on day 0, 7, 14, and 28 after ischemic injury. The ratio of ischetnic/non-ischemic perfusion was quantitatively analyzed (n=6 mice/group). (B-C) CD31 expression in ischemic hind limb by Western blotting. β-actin was used as loading control. (D-E) Anti-CD31 immunostaining in gastrocnemius muscle on day 28 after HLI. showing CD3 1-positive capillaries per muscle fiber with dystrophin staining (n=6 mice/group. Scale bar: 100 μm). (F) EPC (shown as Sca-l^+^/Flk-l+cells) mobilization after tissue ischemia in T1DM mice after administration of GLP-1 (32-36) was determined by flow cytometry. (G) Percentage of Sca-1^+^/Flk-1^+^ cells from different groups (mean ± SEM from 6 individual mice per group). Mean ± SEM from independent experiments. Statistical significance was determined by 1-way ANO VA with post hoc Tukey multiple comparisons test. *P<0.05; **P<0.01; ***P<0.001 ; ns, no significance.

### 2.2 GLP-1(32-36) protects mitochondria from high glucose-induced damage by enhancing glycolytic metabolism

Endothelial metabolism plays an important role in regulating angiogenesis and mitochondrial membrane remodeling is highly responsive to changes in cell metabolism(29). As EPCs are considered to control the angiogenic switch of many physiological and pathologic processes, such as neovascularization (30), we assessed whether GLP-1(32-36) plays a role in improving angiogenesis by regulating mitochondrial dynamics and metabolism. Mice primary bone marrow EPCs (mEPCs) from T1DM mice or T1DM mice injected with GLP-1(32-36) were isolated and cultured, and the 3-to 5-passage mEPCs were used for further experiments. (Supplementary Figure 3). Ultrastructure examination revealed significant increase in elongated mitochondria of mEPCs from STZ-induced diabetic mice treated with GLP-1(32-36), whereas mitochondria from diabetic mice were mostly round or circular (Fig. 3A, B). HG-stimulated accumulation of mitochondrial ROS and loss of mtDNA content were suppressed by GLP-1(32-36) treatment (Fig. 3C, D). GLP-1(32-36) also reversed the effect of HG on mitochondrial membrane potential as well as basal and maximal oxygen consumption rate (OCR) in EPCs from T1DM mice (Fig. 3E-H).

**Figure 3.**
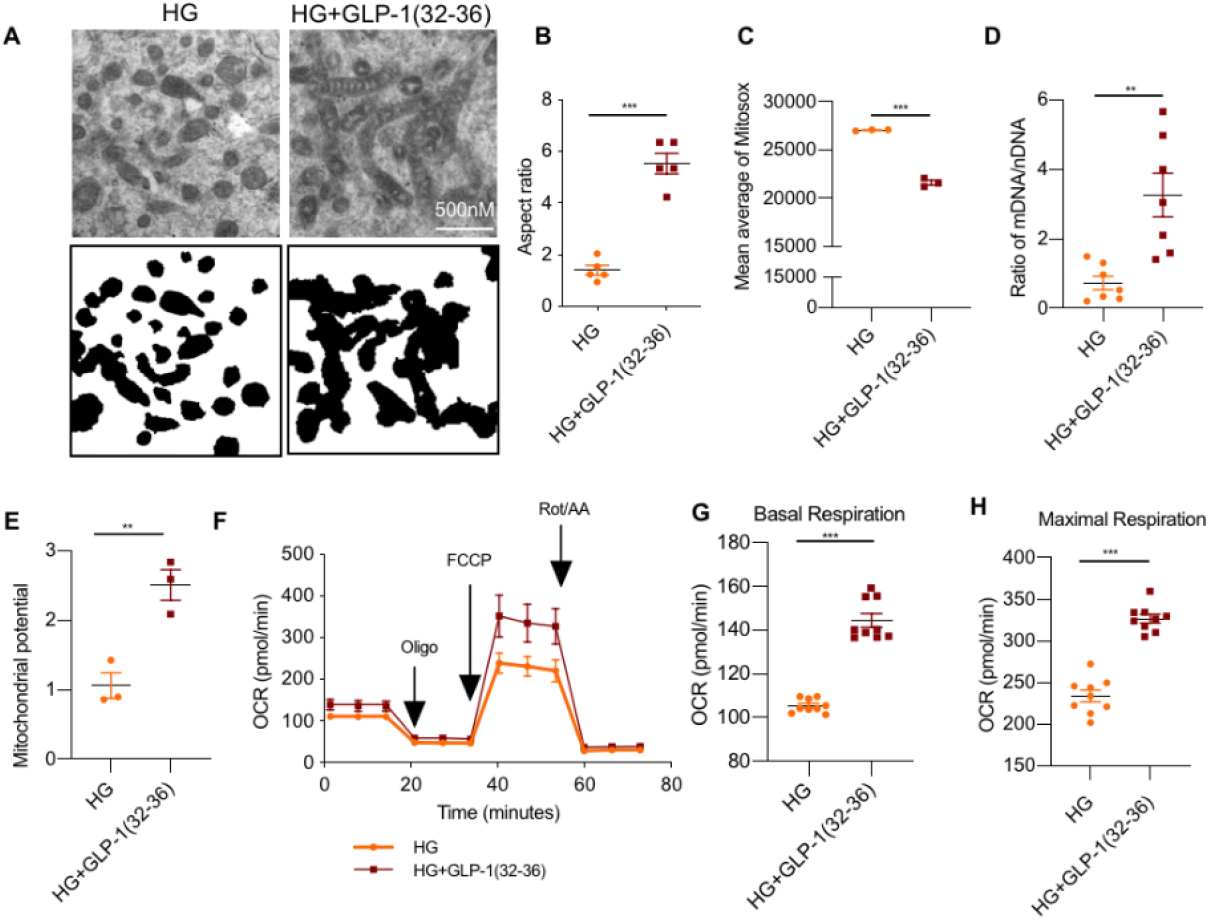
GLP-l(32-36) rescues mitochondrial dynamics and function of mEPCs. Mice primary hone marrow EPCs (mEPCs) from T1DM mice or T1DM mice injected with GLP-l(32-36) were isolated and cultured, and the 3-to 5-passage mEPCs were used for further experiments. (A) Representative TEM micrographs of EPCs mitochondria (top panel). Tracing of mitochondria from TEM micrographs (bottom panel). (B) Average aspect ratio for each group from A. (C) Mitochondrial ROS shown as fluorescence intensity of MitoSOX measured by flow cytometry. (D) mtDNA copy numbers relative to nuclear DNA (nDNA) detected by RT-PCR. (E) Mitochondrial membrancepotential shown as JC-1 fluorescence ratio between red and green. (F) Sequential oxygen consumption rate (OCR) curve was assessed using the Seahorse XF96 analyzer. (G-H) Representative basal and maximal respiration are shown, Quantification of OCR was calculated. Mean ± SEM from independent experiments (B, n=5-7; C, n=3; D, n=7; E, n=3; G&H, n=9) Statistical significance was determined by 1 -way ANOVA with post hoc Tukey multiple comparisons test. *P<(l.05; **P<0.01; ***P<0.001.

By metabolomic analysis, we observed that GLP-1(32-36) treatment induced a compensatory increase in glycolysis in mEPCs from T1DM mice as demonstrated by increased glycolytic metabolites such as L-lactate, D-fructose 1,6-bisphosphate (FDP), dihydroxyacetone phosphate (DHAP), and phosphoenolpyruvate (PEP) (Fig. 4A-B). Eight of the ten genes involved in the aerobic glycolytic pathway were significantly up-regulated by GLP-1(32-36) (Supplementary Table 2, Fig. 4C), including PFKFB3 (6-phosphofructo-2-kinase/fructose-2,6-bisphosphatase 3) which is engaged in the rate-limiting step in glycolysis for increased pyruvate production. The extra pyruvate generated could be directed to lactate rather than to the TCA cycle by upregulation of Lactate dehydrogenase A (LDHA) (Fig. 4D). Seahorse assay also confirmed that GLP-1(32-36) enhanced glycolytic flux (extracellular acidification rate [ECAR]) (Fig. 4E-F). These data demonstrate that GLP-1(32-36) may ameliorate HG-induced excessive mitochondrial fission by resorting to aerobic glycolysis.

**Figure 4.**
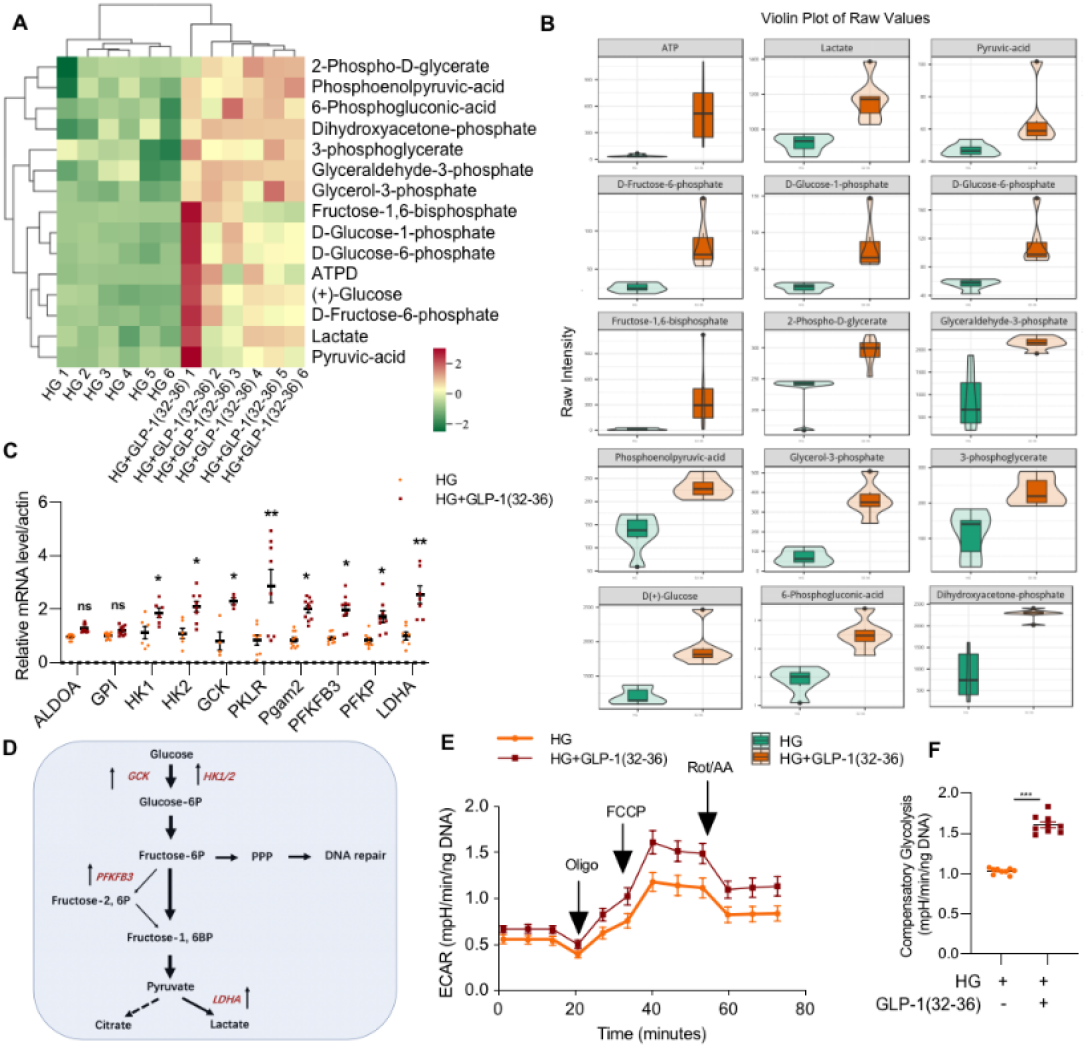
GLP-l(32-36) facilitates glucose flux through glycolysis and transcription of glycolysis-related genes. Metabolome analysis of EPC extracted from mouse marrow (n=6). The cells were divided into two groups with corresponding treatment (HG vs HG+GLP-1 (32-36)). (A) Heatmap of glycolytic related metabolites across all experimental animals of both groups. Red represents high levels and green, low levels. (B) Violin plotting to show data distribution and its probability density, with the box in the middle representing the quartile range; the thin black line extending from it, the 95% confidence interval; the black horizontal line in the middle, the median; and the outer shape, the distribution density of the data. (C) mRNA changes were validated by qPCR of selected glycolytic genes. (D) Upregulation of selected genes within the glycolytic pathway in EPCs from T1DM mice receiving GLP-1 (32-36). (E-F) Glycolytic flux (ECAR, extracellular acidification rate using Seahorse XF) in EPCs from two group of TIDM mice (n = 9/time point). Data in C, E and F represent mean ± SEM from independent experiments (C & F; n=6). Statistical significance was determined by Student’s 2-tailed t test. *P<0.05; **P<0.01; ***P<0.00l; ns, no significance.

### 2.3 GLP-1(32-36) improves angiogenesis via PFKFB3-mediated glycolysis

As PFKFB3 is a well-known glycolytic activator(31), we explored whether PFKFB3 is required for the therapeutic function of GLP-1(32-36) in preventing EPCs angiogenesis disorder. mEPCs were treated with GLP-1(32-36),followed with the PFKFB3-specific inhibitor PFK15 (32) for 30 min. GLP-1(32-36) treatment remarkably upregulated PFKFB3 protein levels and the phosphorylated form of eNOS, which was suppressed by PFK15 treatment (Fig. 5A-C). PFK15 treatment also suppressed GLP-1(32-36)-induced NO secretion, tube formation, and migration area of EPCs (Fig. 5D-F). Taken together, these findings suggest that GLP-1(32-36) recuperates angiogenesis by activating PFKFB3-mediated glycolysis.

**Figure 5.**
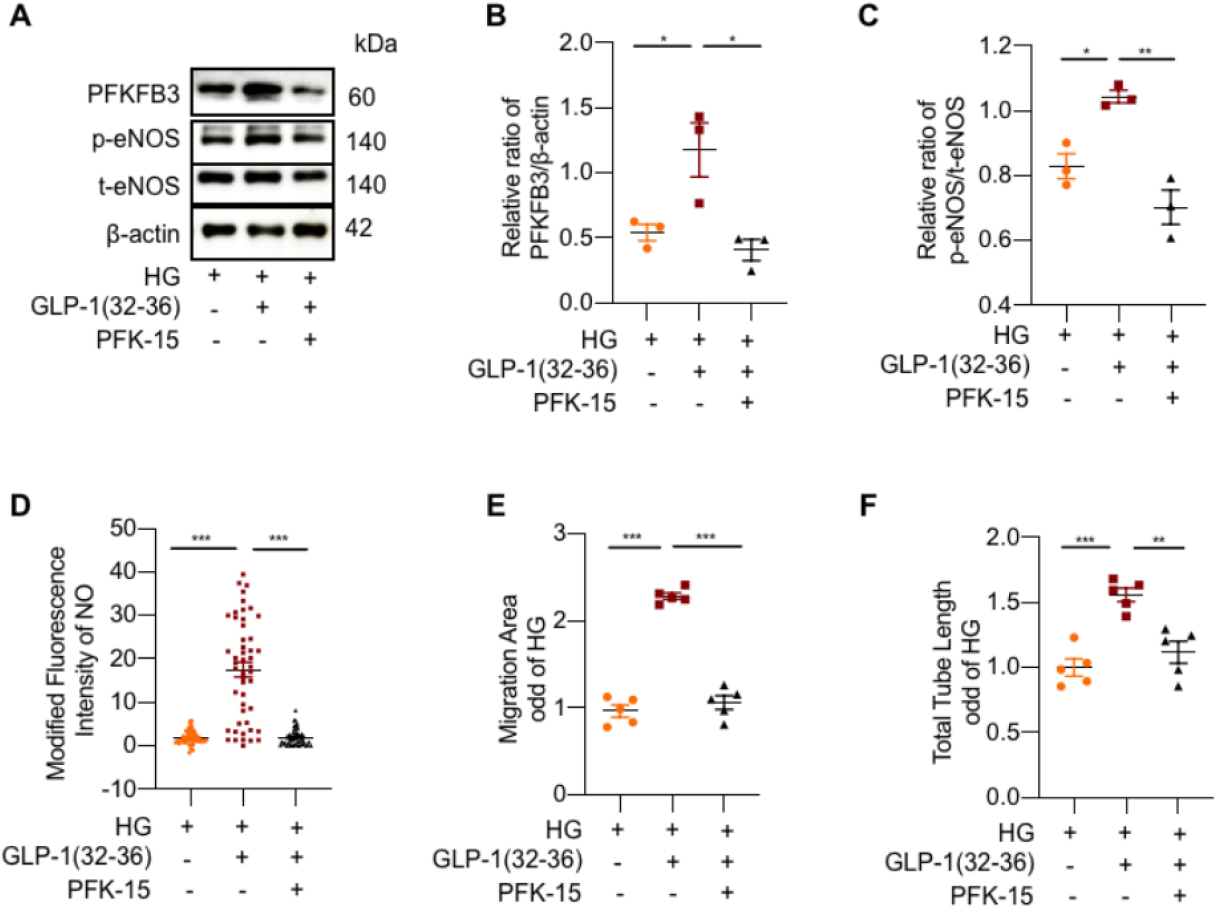
PFKFB3, a key molecule of glycolysis activator, is involved in regulating GLP-l(32-36)-mediated angiogenesis. mEPCs isolated from T1DM mice or T1DM mice injected with GLP-K32-36) were subjected to additional treatment with PFK15, PFKFB3 inhibitor. (A) EPCs extracts were fractionated by SDS-PAGE and analyzed by Western blotting with antibodies to p-eNOS, eNOS and PFKFB3. (3-actin was used as protein loading control. (B) Ratio of p-eNOS to eNOS. (C) Ratio of PFKFB3 to f -actin. (D) The production of NO in mEPCs was measured with a DAF-FM diacetate kit. (E) Quantification of the migration area were taken in 5 random microscopy fields per sample. (F) Quantification of the total tube length in tube formation assay. Data were shown as mean ± SEM from independent experiments (B & C, n=8; D, n=40-50; E & F; n=5f Statistical significance was determined by 1-way ANOVA with post hoc Tukey multiple comparisons test. *P<0.05; **P<0.01; ***P<0.001.

### 2.4 GLP-1R is required for angiogenetic function of GLP-1(32-36)

To test whether GLP-1R deletion interrupts GLP-1(32-36)-induced neovascularization in T1DM, we established Glp1r knockout mice (*Glp1r*^*-/-*^ mice) by the CRISPR/Cas9 technology (Supplementary Fig 4, 5, Fig. 6A). The *Glp1r*^*-/-*^ mice were allocated into four groups and one group infused with 1 x 10^6^ mBM-EPCs overexpressing GLP-1R (Ad-GLP-1R) or the Lv-NC control lentiviral vector via tail vein injection (Supplementary Fig 6). GLP-1(32-36) treatment had marginal effect on blood perfusion in the *Glp1r*^*-/-*^ mice and those receiving control lentivirus (EPCs/Lv-NC), but time-dependently increased blood perfusion in mice received EPCs/Ad-GLP-1R from day 3 to 21 after transplantation (Fig. 6B-C). The benefit of EPC/Ad-GLP-1R was demonstrated by increased expression of CD31, an endothelial cell marker, in ischemic gastrocnemius muscle measured 28 days after HLI (Fig. 6D-E). These findings indicate that GLP-1(32-36)-mediated angiogenesis in T1DM mice after HLI is dependent on GLP-1R expression.

**Figure 6.**
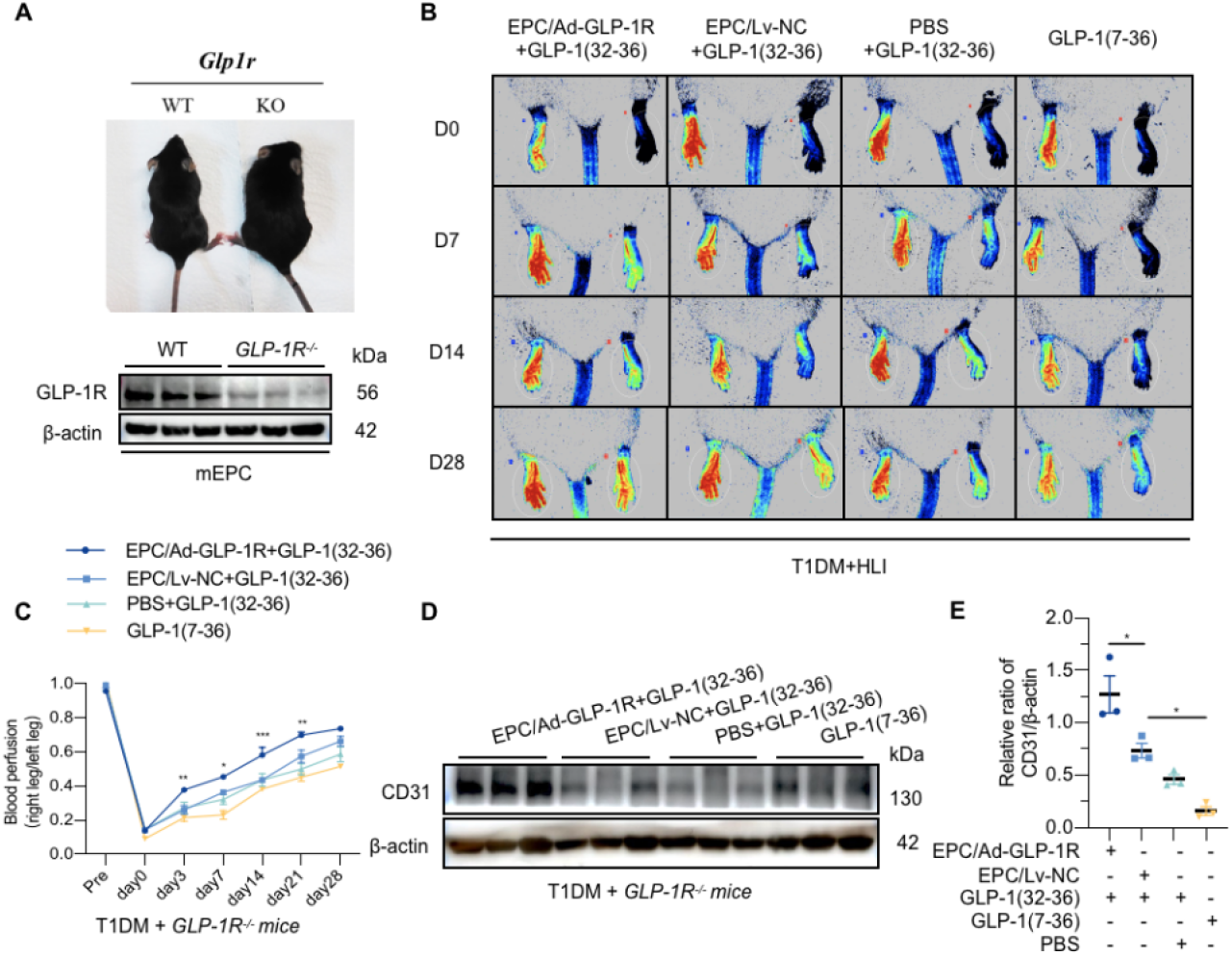
GLP-l(32-36) depends on the cognate GLP-1 R to exert its role in improved blood perfusion and angiogenesis of ischemic limb of T1DMmice. (A) Mating strategy to generate GLP-1 R knockout in diabetic mice and western blotting detected the expression of GLP-IR in WT and KO mice. (B) Representative images showing blood flow reperfusion assessed by Doppler laser ultrasound on day 0, 7, 14, and 28 after ischemic injury. (C) Time courses of blood perfusion shown in images and quantitative analysis after HLI surgery of GLP-IR^−/−^-mice with or without GLP-1 R-expressing EPC transplantation in response to GLP-l (32-36) treatment. (D) Western blotting of CD31 expression in gastrocnemius muscle from GLP-1 R^−/−^-mice receiving the cell therapy on day 28. β-actin was used as a loading control. The gels shown here are representative of three individual experiments. (E) Ratio ofCD31 to βactin. Data represent mean ± SEM from independent experiments (B-H, n=6-8; J, n=3). Statistical significance was determined by 1-way ANO VA with post hoc Tukey multiple comparisons test. *P<0.05; **P<0.01; *** P<0.001 ; ns, no significance.

### 2.5 GLP-1R binds to, but is not required for, GLP-1(32-36) entry into endothelial progenitor cells

To address the question whether GLP-1(32-36) undergoes cellular uptake via GLP-1R, a Cy5-tagged GLP-1(32-36) probe (Cy5-GLP-1(32-36)) was used for fluorescent tracing (Supplementary Table 1, Supplementary Fig 7). Confocal imaging showed that Cy5-GLP-1(32-36) entered EPCs with the strongest intracellular signals observed within 30 min (Supplementary Fig 8). Interestingly, while treating the cells with endocytosis inhibitor Dyngo-4A almost completely blocked abrogate Cy5-GLP-1(32-36) internalization, deletion of GLP-1R had no significant effect on Cy5-GLP-1(32-36) entry into the cells, implicating that GLP-1(32-36) penetration is independent of GLP-1R (Fig. 7A-B).

**Figure 7.**
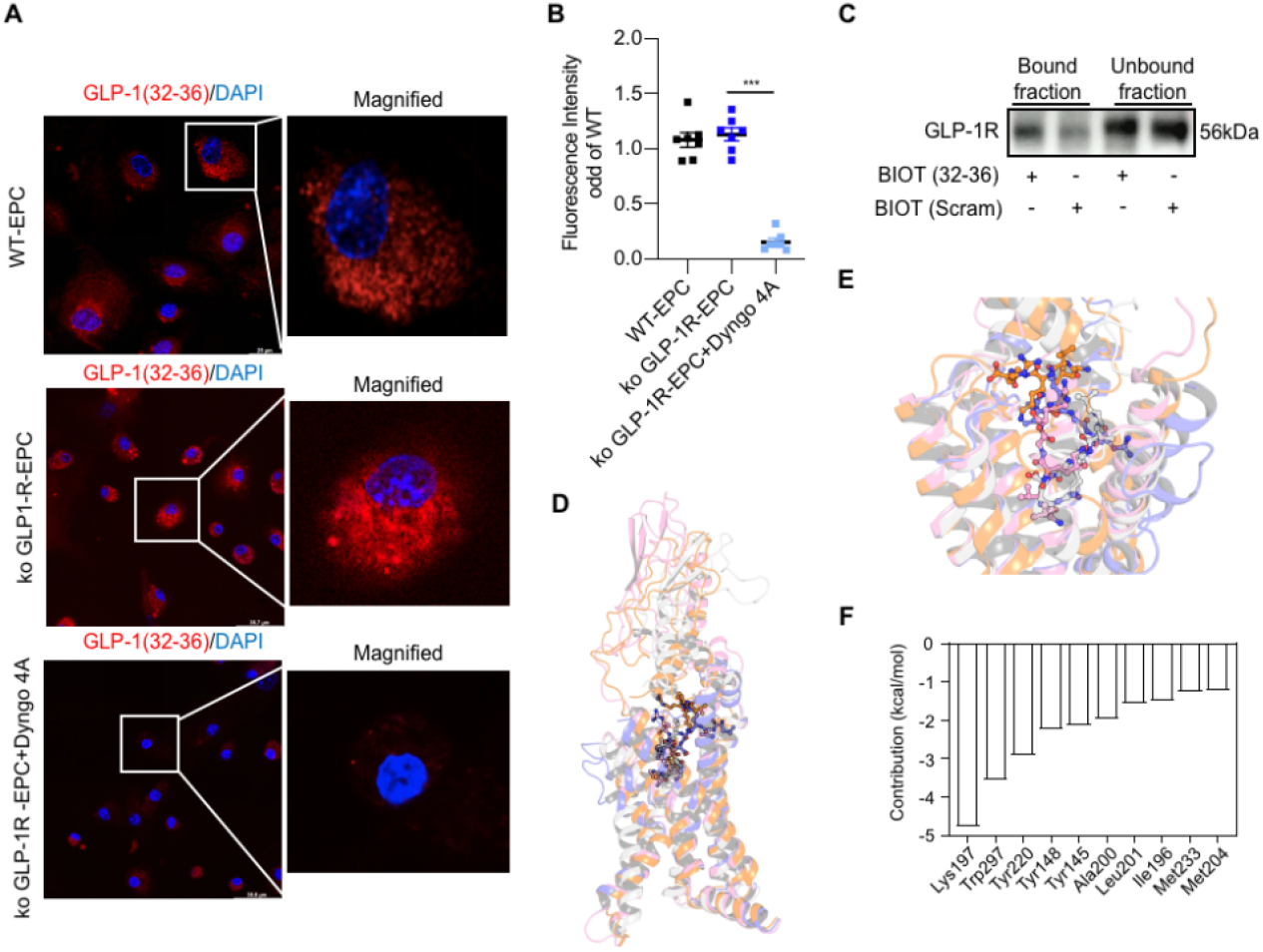
GLP-l(32-36) binds to GLP-1R without requiring its mediation for entry. (A) mEPCs isolated from wild-type and Glplr-/-mice were incubated with 100μM Cy5 tagged GLP-1 (32-36) with or without dynamin inhibitor Dyngo4A (30μM) for 30 (as above) min. and then imaged with spinning disk confocal microscope. (B) Fluorescent intensity of Cy5-GLP-1(32-36) was quantified using Imaged software. (C) Western blotting analysis of GLP-1R in bound and unbound fractions from pull-down experiments showed preferential binding of GLP-1R to biotinylated GLP-1 (32-36) [BIOT (32-36)] as compared with the biotinylated scrambled (32-36) control [BIOT (scram)]. (D) Prediction of the binding mode between GLP-1(32-36) and GLP-1R. Overview of global docking of peptide to 4 crystal structures of GLP-1 R (5NX2: light gray, 60RV: light blue, 6X18: pink, 7C2E: orange). Detailed view of global docking of GLP-1 (32-36) to 4 crystal structures of GLP-1 R (5NX2: light gray, 6ORV: light blue, 6X18: pink, 7C2E: orange). (F) Top-ranking key residues of GLP-1R in binding with GLP-l(32-36). Data were mean ± SEMfrom independent experiments (B, n=8). Statistical significance was determined by I-way ANOTA with post hoc Tukey multiple comparisons test. *P<0.05; **P<0.01; ***P<0.001.

To determine binding to GLP-1R is necessary for the angiogenetic effect of GLP-1(32-36), we conducted an affinity pull-down experiment with biotinylated GLP-1(32-36) (BIOT (32-36)) (Supplementary Table 3). In agreement with the finding of others (33), we found that the pentapeptide bound to GLP-1R (Fig. 7C). Global docking analysis revealed that GLP-1(32-36) bound to the GLP-1 binding site of GLP-1R for all crystal structures (Figure 7D-F). These data suggest that interaction with GLP-1R, though is not involved in GLP-1(32-36) entry into the cells, is required for activation of GLP-1R-mediated pathway to play its subsequent angiogenetic effect.

### 2.6 GLP-1(32-36) exerts its angiogenetic effect through the eNOS/cGMP/PKG pathway via GLP-1R

Human EPCs (hEPCs) were isolated from cord blood for its richness in EPCs (34), cells were isolated and identified by specific EPCs markers as previously described (35) (Supplementary Fig. 9). We found that GLP-1(32-36) increased generation of cGMP but not cAMP (Supplementary Fig 10), indicating that the pentapeptide might activate the eNOS/cGMP/PKG pathway to exert angiogenesis. Downregulation of GLP-1R by siRNA silencing attenuated GLP-1(32-36)-induced cGMP increase and NO formation (Supplementary Fig. 6B; Fig. 8A-B). Suppressing GLP-1R also recued GLP-1(32-36)-stimulated p-eNOS and PKG expression (Fig. 8C-E). GLP-1(32-36) treatment resecured HG-induced impairement in tube formation and migration, but this effect was suppressed by GLP-1R downregulation (Fig. 8E-F). These data uncover a mechanism whereby GLP-1(32-36) improves angiogenesis in HG-exposed EPCs.

**Figure 8.**
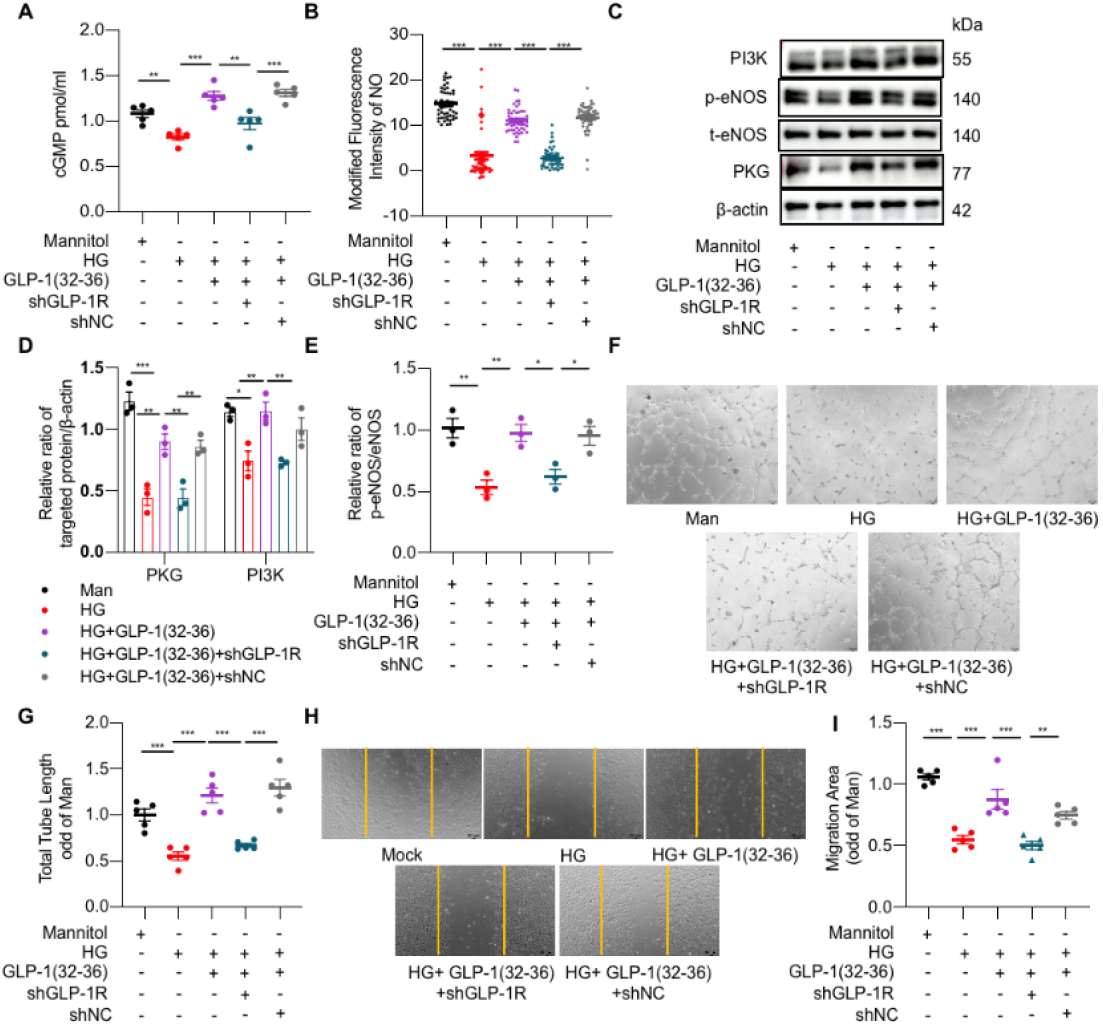
Both GLP-l(32-36) and GLP-1 R are required in improving angiogenesis via the eNOS/cGMP/PKGpathway. hEPCs were infected with lentivirus expressing shRNA targeting Glplr for 12 h and treated with GLP-1 (32-36) (100 nM). and then incubated in high glucose (33 mM) for 24 h. Mannitol (Man) was used as the control. (A) cyclic GMP was detected by using the cGMP ELISA Kit (n=5). (B) NO secretion was measured with a diacetate diaminofluorescein-FM (DAF-FM) diacetate kit and was imaged on a fluorescence microscope. (C) EPCs extracts were fractionated by SDS-PAGE and analyzed by Western blotting with antibodies against p-eNOS (Seri 177) and eNOS, PI3K and PKG.β -actin was used as loading control. The gels shown here are representative of 3 individual experiments. (D) Ratio of PKG or PI3K to fi-actin (E) Ratio of p-eNOS to eNOS. (F-G) Quantification of the total tube length in EPCs, images of tube morphology were taken in 5 random microscopy fields per sample. (H-I) Quantification of the migration area were taken in 5 random microscopy fields per sample. Scale bars: 500 μm. Data were mean ± SEM from independent experiments (A, n=5; B, n=50; D&E, n=3; F&G, n=5). Statistical significance was determined by 1-way ANOTA with post hoc Titkey multiple comparisons test. *P<0.05; **P<0.01; ***P<0.001.

### 2.7 GLP-1R is involved in mitochondrial dynamics and GLP-1(32-36)-mediated glycolysis

We examined the potential effect of GLP-1(32-36) on mitochondrial biogenesis in EPCs treated the GLP-1R antagonist (exendin (9-39)). GLP-1(32-36) recovered the mitochondrial morphology, which is blocked by exendin (Supplementary. 11A-E). Inhibition of GLP-1R also interrupted the improvement of MMP and mitochondrial OCR level by pentapeptide (Supplementary. 11F-I). These results suggest that GLP-1R is involved in pentapeptide-mediated mitochondrial homeostasis.

Given that PFKFB3 plays an important role in vessel sprouting by promoting glycolysis(31), we wondered if GLP-1R is involved in GLP-1(32-36)-mediated effects on metabolism. Suppressing GLP-1R by shRNA impaired ATP turnover, abolished the positive effect of GLP-1(32-36) on lactate accumulation (Fig. 9A-B), and blocked the effect of GLP-1(32-36) on glucose uptake (Fig. 9C-D). More importantly, downregulation of GLP-1R suppressed PFKFB3 expression (Fig. 9E-F). These data reveal that GLP-1R is responsible for GLP-1(32-36)-mediated glycolysis via PFKFB3.

**Figure 9.**
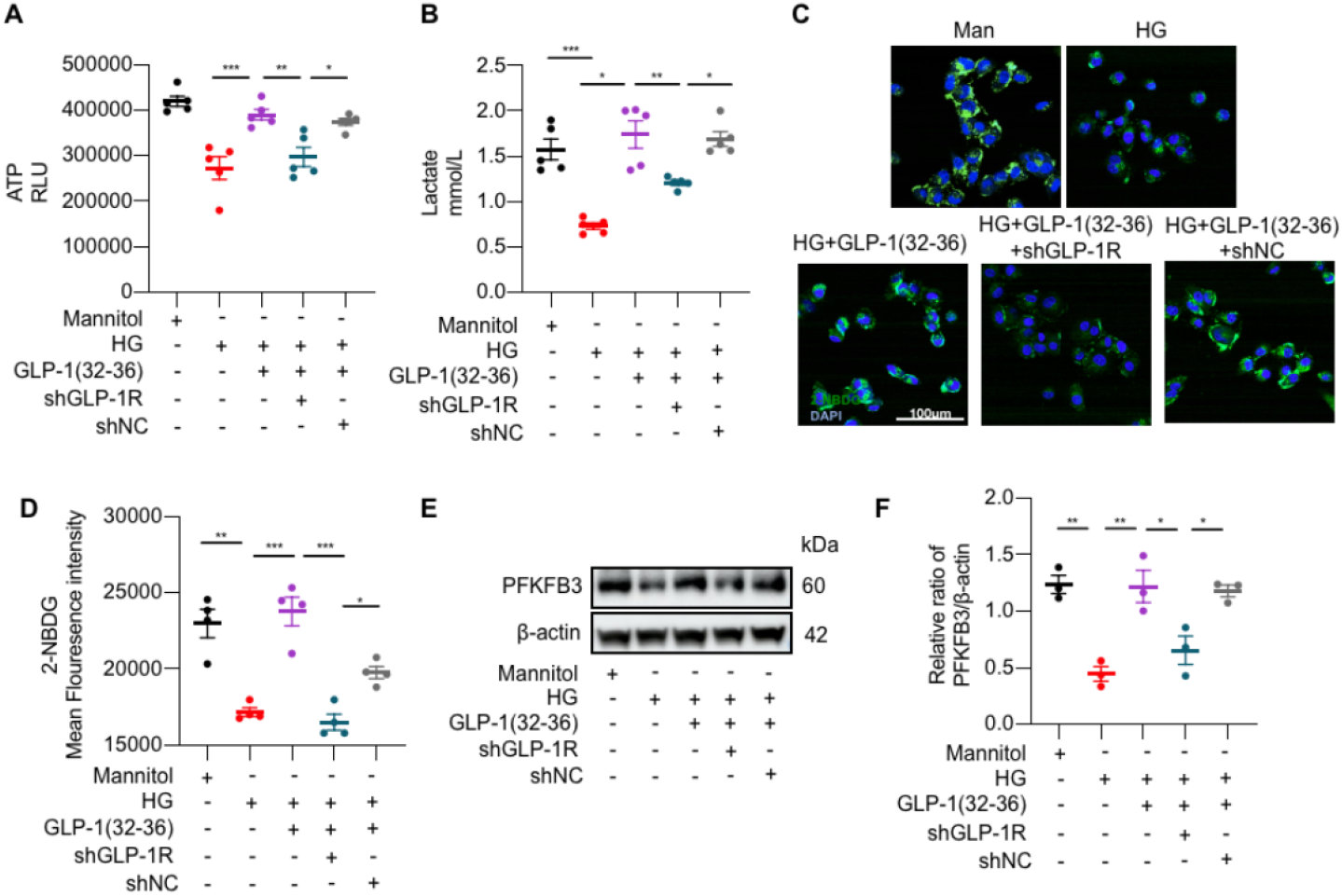
Both GLP-1 (32-36) and GLP-1R are involved in regulating PFKFB3-mediated glycolysis. hEPCs were infected with the lentivirus expressing shRNA targeting Glplr for 12 h and treated with GLP-1 (32-36) (100 nM), and then incubated in high glucose (HG, 33 mM) for 24 h. Mannitol (Man) was used as the control. (A) ATP content. (B) Lactate production. (C-D) Uptake of 2-NBDG as determined by immunofluorescence staining and flow cytometry analysis. (E) EPCs extracts were fractionated by SDS-PAGE and analyzed by Western blotting with PFKFB3. β-actin was used as loading control. (F) Ratio of PFKFB3 to β-actin. The gels shown here are representative of 3 individual experiments. Data are expressed as mean ± SEMfrom independent experiments (A, n=5; B, n=6; D, n=4; F, n=3). Statistical significance was determined by 1-way ANOVA with post hoc Tukey multiple comparisons test. *P<0.05; **P<0.01; ***P<0.001.

## 5. Discussion

GLP-1(9-36) is known to improve hypoxia-impaired human aortic endothelial cell viability (11) and attenuate HG-triggered mitochondrial ROS generation in human endothelial cells (12). Of the two peptides cleaved from GLP-1(9-36) amide by NEP 24.11(10), GLP-1(28-36) could activate the AC-cAMP signaling pathway and play a role in cardiovascular protection (13) and GLP-1 (32-36) may increase energy consumption, decrease body weight and decrease β-cell apoptosis in obese mice (14, 15). However, if and how GLP-1(32-36) contributes to protection of vascular endothelial injury remains unexplored. Using in vivo, ex vivo and in vitro models, we demonstrate that GLP-1(32-36) is effective in rescuing angiogenic function, blood perfusion and promoted EPC mobilization in ischemic limb of STZ-induced diabetic mice independent of insulin release. Mechanistically, it recuperates angiogenesis by activating PFKFB3-mediated aerobic glycolysis and ameliorate excessive mitochondrial fission. Its interaction with GLP-1R is required for regulation of the glycolytic effect via PFKFB3 as well as for modulation of the eNOS/cGMP/PKG pathway to improve angiogenesis in high glucose-exposed EPCs. This study explores the mechanism by which GLP-1(32-36) promotes angiogenesis and establishes the theoretical basis for the clinical development and application of angiogenic agents in diabetic foot patients.

PFKFB3 plays important roles in angiogenesis in several cell types (31). In cancer-associated fibroblasts and tumor endothelial cells, PFKFB3 is known to modulate angiogenesis via activation of the aerobic glycolysis, and blockade of PFKFB3 reduces tumor angiogenesis(36), while its expression is increased markedly in beta cells from type 1 diabetes patients as well as in human and rat islets exposed to cytokines(37), and in kidneys from diabetic mice(38). Transcriptionally, PFKFB3 expression is regulated negatively by PGC1α(39, 40) or Kruppel-like factor 2 (KLF2)(41), but positively by HIF-1α in response to hypoxia(42, 43) . HIF1α itself could be activated by ROS (44) or endothelial cell-sourced NO in astrocytes(45). Because GLP-1(32-36) increased PFKFB3 without affecting cAMP in HG-exposed EPCs, and PGC1α negatively regulates PFKFB3, it is less likely that the pentapeptide acts via the GLP-1R/cAMP/AMPK/PGC1α pathway(40, 46). Activation of HIF1α-PFKFB3 axis is believed to divert glucose metabolism in diabetic β-cells away from mitochondria for glycolysis(44). Therefore, we propose that GLP-1(32-36) exerts its pro-angiogenetic effect in the EPCs exposed to high glucose by activating PI3K/eNOS/cGMP/PKG pathway with enhanced glycolysis and improved mitochondrial homeostasis via PFKFB3 upregulation. This is based on the facts that GLP-1(32-36) could derepress a series of molecules PI3K, NO/eNOS, cGMP and PKG which are involved in myocardial ischemic preconditioning(47), and increase expression of PFKFB3 possibly via NO activation of HIF1α (44) leading to glucometabolic reprogramming shown as diversion of pyruvate away from mitochondria(40, 43).

GLP-1 is known to exert its effect through the unique GLP-1R in stimulating adenylate cyclase activity, and the resultant accumulation cyclic AMP (cAMP) leads to activation of protein kinase A (PKA), one of the multiple intracellular mediators in various tissue (48, 49). According to the ‘‘dual receptor theory’’ (16), GLP-1 may exert insulin mimetic actions on insulin-sensitive target tissues either by acting on GLP-1R to activate the pro-survival cAMP/PKA and PI3K/Akt pathways (49) or through an alternative mechanism independent of GLP-1R as seen with GLP-1(9-36) through a novel receptor/transporter(11, 16, 18). However, whether GLP-1R is involved in cellular uptake and activities of GLP-1(32-36), a further cleavage product derived from GLP-1(7-36) and GLP-1(9-36), has not been determined. We show that cellular entry of GLP-1(32-36) was not affected by silencing of GLP-1R, but completely blocked by the dynamin inhibitor, suggesting that the pentapeptide enters the cell via endocytosis, as with most of the peptide hormones (33). By affinity pull-down and molecular docking analysis, we found that GLP-1R, though not required for GLP-1(32-36) uptake, might act as the binding partner.

Endothelial dysfunction in association with cGMP-NO insufficiency has been well established as an important correlate of heightened cardiovascular risk (50). NO is short-lived and acts in autocrine and paracrine manners by elevating cGMP/PKG, thereby exerting cardioprotective effects on remodeling(51). NO synthase null mutant mice displayed markedly reduced mitochondrial content associated with significantly lower oxygen consumption and ATP content(52). We hypothesized that GLP-1(32-36) might mediate eNOS/cGMP/PKG-dependent mitochondrial biogenesis to promote angiogenesis via GLP-1R. In our study, GLP-1(32-36) stimulated cGMP (but not cAMP) production, promoted NO release, eNOS phosphorylation and PKG expression, and enhanced tube formation and migration, all of which were suppressed by downregulation of GLP-1R. These data support a mechanism whereby GLP-1(32-36) utilizes GLP-1R to modulate the eNOS/cGMP/PKG pathway, independent of G-protein signaling, to play its roles in angiogenesis and glycolysis.

In addition, eNOS-derived NO is well known to tightly regulate mitochondrial functioning. In physiological concentrations, NO regulates mitochondrial network fusion by phosphorylating and inhibiting dynamin related GTPase (DRP1) through the sGC/PKG pathway(53). Our data also define the molecular mechanism of mitochondrial dynamics and metabolism mediated by GLP-1(32-36) through eNOS/cGMP/PKG pathway. Previous studies have showed that GLP-1 byproducts could target mitochondria upon entry into the cells. GLP-1(9-36) could reduce elevated levels of mitochondrial-derived ROS in Alzheimer’s disease model mice(54). GLP-1(28-36) prevents ischemic cardiac injury by inhibiting mitochondrial trifunctional protein-α(13). It has been proposed that its C-terminal domain, VKGR amide, might contain a consensus mitochondrial targeting sequence(55). This suggests that GLP-1(32-36) might also play a role in regulating mitochondria fitness. We constructed Cy5-conjugated pentapeptide to visualize its internalization. Cy5-GLP-1(32-36) entered EPCs with the strongest intracellular signals observed within 30 min. Pharmacological inhibition of GLP-1R by chemical antagonists (exendin (9-39)) blunted peptides-regulated mitochondrial homeostasis and dynamics by tilting mitochondrial fission towards fusion. We found that GLP-1(32-36) rescues mitochondrial morphology and protected oxidative stress injury from high glucose stress. These findings provide evidence that GLP-1(32-36) is involved in improvement of mitochondrial fitness.

Since endothelial function is intrinsically linked to its metabolism(56, 57). ECs are atypical nonmalignant cells that surprisingly depend on glycolysis to synthesize>80% of ATP even under well-oxygenated conditions(31, 58). We hypothesized that occurrence of PAD could be accompanied with metabolic abnormalities and failure to establish timely metabolic switch might negatively affect the function of endothelial cells. We investigated the metabolite profiles between mEPCs from T1DM with GLP-1(32-36) treatment and from control T1DM (without treatment). The results showed that high glucose stress resulted in glycolysis disorder which was rescued by GLP-1(32-36). Seahorse data also show that GLP-1(32-36) could improve basal and maximal respiration and glycolysis in mEPCs exposed to high glucose. Transcription of key genes involved in the aerobic glycolytic pathway and lactate level were increased by GLP-1(32-36) treatment. The elongated mature mitochondria of GLP-1(32-36)-treated EPCs and increased glycolysis, as opposed to immature round mitochondria lacking mature cristae characteristic of hyperglycemia, illustrates the importance of efficient biosynthetic organelles and efficient energy metabolism to support EPC proliferation required for angiogenesis.

Interestingly, we compared insulin secretion after treating GLP-1(7-36) and GLP-1(32-36) with different concentration. GLP-1(7-36) stimulates glucose-dependent insulin secretion, whereas GLP-1(32-36) has only weak partial insulinotropic agonist activities. This not only elucidates the key role of GLP-1R in the action of GLP-1(32-36) but also suggested GLP-1(32-36) has self-governed angiogenesis ability independent of insulin release and blood glucose control, expanding the possibility of the pentapeptide improving angiogenesis in non-diabetic patients.

In summary, we have found that GLP-1(32-36) functions in cooperation with GLP-1R in mediating eNOS-cGMP-PKG signaling, mitochondrial homeostasis and metabolic functions of endothelial cells in favor of angiogenesis. GLP-1(32-36) prevents HG-mediated mitochondrial fission, regulates metabolic reprogramming by enhanced PFKFB3 expression and glycolysis, and improve EPCs angiogenesis. Our results are consistent with an emerging appreciation that GLP-1R mediated mitochondrial dynamics and energy metabolism plays a central role in endothelial angiogenesis. We believe that the mechanistic pathway uncovered here has potential impact, from the clinical perspective, on therapy of PAD, for which no approved therapies currently exist. GLP-1(32-36) may offer supplementary protection against ischemic angiogenesis in instances where reduced blood flow-related mitochondrial fitness and glycolysis is compromised before apparent clinical manifestation.

## 4. Materials and Methods

The origins and specifications of the mice used in this study are detailed in the Supplemental Methods. Descriptions of detailed methods of T1DM in vivo (26) and mouse HLI models in vivo (59) are provided in the Supplemental Methods. Details on the isolation of mEPCs and hEPCs, and cell culture methods, including drug treatment regimens, are provided in the Supplemental Methods. All assays relevant to cAMP, cGMP, and Seahorse XFe96 extracellular flux measurements are detailed. Gene silencing using siRNA and Western blotting protocols are described in the Supplemental Information.

### 4.1 Statistics

Data are presented as mean ± SEM. Comparisons between two groups were performed by Student’s 2-tailed t test. Comparisons of data with three or more groups were performed using 1-way ANOVA with Tukey’s post hoc multiple comparisons. Repeated-measures ANOVA was performed when appropriate. All statistical analyses were performed in GraphPad Prism (Version 9.0). In all cases, differences were considered significant at *P < 0.05 and highly significant at **P < 0.01 and ***P < 0.001; ns means no significance.

### 4.2 Study approval

Experimental setups and animal care were permitted by the Animal Policy and Welfare Committee of Zhejiang University (ethical approval code:2021-NO.172). And for human studies, the work was approved by the Ethics Committee of Zhejiang University (ethical approval code: 2021-NO.0318) and carried out in accordance with the Declaration of Helsinki. All participants gave written informed consent.

## Author Contributions

Yikai Zhang†: Conceptualization, designing research studies and writing the manuscript, Shengyao Wang†: Conducting experiments, acquiring date and analyzing data. They shared first authorship. Qiao Zhou, Yepeng Hu and Yi Xie: Conducting experiments and analyzing data. Weihuan Fang: reviewing & editing the manuscript. Zhe Wang, Shu Ye and Xinyi Wang: Data curation. Chao Zheng*: Supervision, administrating project and funding the study.

## Funding

We would like to express our thanks to the Natural Science Foundation of China (NO.82100862, NO.82070833), Zhejiang Provincial Natural Science Foundation (NO. LY22H070001, NO. LZ19H020001), and Zhejiang Provincial Key Research & Development Program (NO.2021C03070) for financial support.

## Conflicts of Interest

The authors declare that there is no conflict of interest regarding the publication of this article.

## Supplementary Materials

**Supplemental methods** contain additional details of *in vitro* and *in vivo* experiments, along with relevant reference citations.

**Fig. S1**. GLP-1(32-36) improves angiogenesis of HUVECs in hyperglycemia.

**Fig. S2**. Streptozotocin-induced diabetic models in Mice.

**Fig. S3**. Characterization of isolated mice bone marrow EPCs.

**Fig. S4**. *GLP1r* Cas9-KO strategy.

**Fig. S5**. Diabetic model in *GLP-1*^*-/-*^ with transplanting EPCs/Lv-GLP-1R or control. **Fig. S6**. Verification the effect of lentivirus overexpression in mBM-EPCs and hUCB-EPCs.

**Fig. S7**. Synthesis of Cy5-GLP-1(32-36).

**Fig. S8**. Cellular uptake of GLP-1(32-36).

**Fig. S9**. Characterization of isolated human EPCs(hEPCs).

**Fig. S10**. GLP-1(32-36) do not stimulate intracellular cAMP accumulation but cGMP in EPCs.

**Table S1**. Amino acid sequences of synthetic peptides used.

**Table S2**. List of primer sequences used for quantitative RT-PCR.

**Table S3**. Amino acid sequences of biotinylated peptides used.

**Table S4**. Lentivirus sequences to inhibit or overexpress gene expression.

## Notes

### Competing Interest Statement

The authors have declared no competing interest.

## References

1. Ungerleider JL, Christman KL. Concise Review: Injectable Biomaterials for the Treatment of Myocardial Infarction and Peripheral Artery Disease: Translational Challenges and Progress. Stem cells translational medicine. 2014;3(9):1090–9.

2. Klein AJ, Ross CB. Endovascular treatment of lower extremity peripheral arterial disease. Trends Cardiovasc Med. 2016;26(6):495–512.

3. Petznick AM, Shubrook JH. Treatment of specific macrovascular beds in patients with diabetes mellitus. Osteopath Med Prim Care. 2010;4:5.

4. Aronis KN, Chamberland JP, Mantzoros CS. GLP-1 promotes angiogenesis in human endothelial cells in a dose-dependent manner, through the Akt, Src and PKC pathways. Metabolism. 2013;62(9):1279–86.

5. Katare R, Riu F, Rowlinson J, Lewis A, Holden R, Meloni M, et al. Perivascular delivery of encapsulated mesenchymal stem cells improves postischemic angiogenesis via paracrine activation of VEGF-A. Arterioscler Thromb Vasc Biol. 2013;33(8):1872–80.

6. Phillips LK, Prins JB. Update on incretin hormones. Ann N Y Acad Sci. 2011;1243:E55–74.

7. Orskov C, Wettergren A, Holst JJ. Biological effects and metabolic rates of glucagonlike peptide-1 7-36 amide and glucagonlike peptide-1 7-37 in healthy subjects are indistinguishable. Diabetes. 1993;42(5):658–61.

8. Kieffer TJ, Habener JF. The glucagon-like peptides. Endocrine reviews. 1999;20(6):876–913.

9. Beinborn M, Worrall CI, McBride EW, Kopin AS. A human glucagon-like peptide-1 receptor polymorphism results in reduced agonist responsiveness. Regulatory peptides. 2005;130(1-2):1–6.

10. Hupe-Sodmann K, McGregor GP, Bridenbaugh R, Göke R, Göke B, Thole H, et al. Characterisation of the processing by human neutral endopeptidase 24.11 of GLP-1(7-36) amide and comparison of the substrate specificity of the enzyme for other glucagon-like peptides. Regul Pept. 1995;58(3):149–56.

11. Ban K, Kim KH, Cho CK, Sauvé M, Diamandis EP, Backx PH, et al. Glucagon-like peptide (GLP)-1(9-36)amide-mediated cytoprotection is blocked by exendin(9-39) yet does not require the known GLP-1 receptor. Endocrinology. 2010;151(4):1520–31.

12. Giacco F, Du X, Carratú A, Gerfen GJ, D’Apolito M, Giardino I, et al. GLP-1 Cleavage Product Reverses Persistent ROS Generation After Transient Hyperglycemia by Disrupting an ROS-Generating Feedback Loop. Diabetes. 2015;64(9):3273–84.

13. Siraj MA, Mundil D, Beca S, Momen A, Shikatani EA, Afroze T, et al. Cardioprotective GLP-1 metabolite prevents ischemic cardiac injury by inhibiting mitochondrial trifunctional protein-α. J Clin Invest. 2020;130(3):1392–404.

14. Tomas E, Stanojevic V, McManus K, Khatri A, Everill P, Bachovchin WW, et al. GLP-1(32-36)amide Pentapeptide Increases Basal Energy Expenditure and Inhibits Weight Gain in Obese Mice. Diabetes. 2015;64(7):2409–19.

15. Sun L, Dai Y, Wang C, Chu Y, Su X, Yang J, et al. Novel Pentapeptide GLP-1 (32-36) Amide Inhibits β-Cell Apoptosis In Vitro and Improves Glucose Disposal in Streptozotocin-Induced Diabetic Mice. Chemical biology & drug design. 2015;86(6):1482–90.

16. Tomas E, Habener JF. Insulin-like actions of glucagon-like peptide-1: a dual receptor hypothesis. Trends Endocrinol Metab. 2010;21(2):59–67.

17. Nikolaidis LA, Elahi D, Shen YT, Shannon RP. Active metabolite of GLP-1 mediates myocardial glucose uptake and improves left ventricular performance in conscious dogs with dilated cardiomyopathy. Am J Physiol Heart Circ Physiol. 2005;289(6):H2401–8.

18. Ban K, Noyan-Ashraf MH, Hoefer J, Bolz SS, Drucker DJ, Husain M. Cardioprotective and vasodilatory actions of glucagon-like peptide 1 receptor are mediated through both glucagon-like peptide 1 receptor-dependent and -independent pathways. Circulation. 2008;117(18):2340–50.

19. Tomas E, Wood JA, Stanojevic V, Habener JF. GLP-1-derived nonapeptide GLP-1(28-36)amide inhibits weight gain and attenuates diabetes and hepatic steatosis in diet-induced obese mice. Regul Pept. 2011;169(1-3):43–8.

20. Elahi D, Angeli FS, Vakilipour A, Carlson OD, Tomas E, Egan JM, et al. GLP-1(32-36)amide, a novel pentapeptide cleavage product of GLP-1, modulates whole body glucose metabolism in dogs. Peptides. 2014;59:20–4.

21. Chen J, Wang D, Wang F, Shi S, Chen Y, Yang B, et al. Exendin-4 inhibits structural remodeling and improves Ca(2+) homeostasis in rats with heart failure via the GLP-1 receptor through the eNOS/cGMP/PKG pathway. Peptides. 2017;90:69–77.

22. Eelen G, de Zeeuw P, Treps L, Harjes U, Wong BW, Carmeliet P. Endothelial Cell Metabolism. Physiol Rev. 2018;98(1):3–58.

23. Li X, Sun X, Carmeliet P. Hallmarks of Endothelial Cell Metabolism in Health and Disease. Cell Metab. 2019;30(3):414–33.

24. Li X, Kumar A, Carmeliet P. Metabolic Pathways Fueling the Endothelial Cell Drive. Annu Rev Physiol. 2019;81:483–503.

25. Zheng J, Xie Y, Ren L, Qi L, Wu L, Pan X, et al. GLP-1 improves the supportive ability of astrocytes to neurons by promoting aerobic glycolysis in Alzheimer’s disease. Mol Metab. 2021;47:101180.

26. Wu KK, Huan Y. Streptozotocin-induced diabetic models in mice and rats. Curr Protoc Pharmacol. 2008;Chapter 5:Unit 5.47.

27. Yu J, Dardik A. A Murine Model of Hind Limb Ischemia to Study Angiogenesis and Arteriogenesis. Methods Mol Biol. 2018;1717:135–43.

28. Yan X, Su Y, Fan X, Chen H, Lu Z, Liu Z, et al. Liraglutide Improves the Angiogenic Capability of EPC and Promotes Ischemic Angiogenesis in Mice under Diabetic Conditions through an Nrf2-Dependent Mechanism. Cells. 2022;11(23).

29. Mishra P, Chan DC. Metabolic regulation of mitochondrial dynamics. J Cell Biol. 2016;212(4):379–87.

30. Gao D, Nolan DJ, Mellick AS, Bambino K, McDonnell K, Mittal V. Endothelial progenitor cells control the angiogenic switch in mouse lung metastasis. Science. 2008;319(5860):195–8.

31. De Bock K, Georgiadou M, Schoors S, Kuchnio A, Wong BW, Cantelmo AR, et al. Role of PFKFB3-driven glycolysis in vessel sprouting. Cell. 2013;154(3):651–63.

32. Zhu W, Ye L, Zhang J, Yu P, Wang H, Ye Z, et al. PFK15, a Small Molecule Inhibitor of PFKFB3, Induces Cell Cycle Arrest, Apoptosis and Inhibits Invasion in Gastric Cancer. PLoS One. 2016;11(9):e0163768.

33. Pandey KN. Endocytosis and Trafficking of Natriuretic Peptide Receptor-A: Potential Role of Short Sequence Motifs. Membranes (Basel). 2015;5(3):253–87.

34. Murohara T, Ikeda H, Duan J, Shintani S, Sasaki K, Eguchi H, et al. Transplanted cord blood-derived endothelial precursor cells augment postnatal neovascularization. The Journal of clinical investigation. 2000;105(11):1527–36.

35. Yan X, Cai S, Xiong X, Sun W, Dai X, Chen S, et al. Chemokine receptor CXCR7 mediates human endothelial progenitor cells survival, angiogenesis, but not proliferation. Journal of cellular biochemistry. 2012;113(4):1437–46.

36. Cantelmo AR, Conradi LC, Brajic A, Goveia J, Kalucka J, Pircher A, et al. Inhibition of the Glycolytic Activator PFKFB3 in Endothelium Induces Tumor Vessel Normalization, Impairs Metastasis, and Improves Chemotherapy. Cancer Cell. 2016;30(6):968–85.

37. Nomoto H, Pei L, Montemurro C, Rosenberger M, Furterer A, Coppola G, et al. Activation of the HIF1α/PFKFB3 stress response pathway in beta cells in type 1 diabetes. Diabetologia. 2020;63(1):149–61.

38. Song C, Wang S, Fu Z, Chi K, Geng X, Liu C, et al. IGFBP5 promotes diabetic kidney disease progression by enhancing PFKFB3-mediated endothelial glycolysis. Cell Death Dis. 2022;13(4):340.

39. Prieto I, Alarcón CR, García-Gómez R, Berdún R, Urgel T, Portero M, et al. Metabolic adaptations in spontaneously immortalized PGC-1α knock-out mouse embryonic fibroblasts increase their oncogenic potential. Redox Biol. 2020;29:101396.

40. Li X, Jiang E, Zhao H, Chen Y, Xu Y, Feng C, et al. Glycometabolic reprogramming-mediated proangiogenic phenotype enhancement of cancer-associated fibroblasts in oral squamous cell carcinoma: role of PGC-1α/PFKFB3 axis. Br J Cancer. 2022;127(3):449–61.

41. Doddaballapur A, Michalik KM, Manavski Y, Lucas T, Houtkooper RH, You X, et al. Laminar shear stress inhibits endothelial cell metabolism via KLF2-mediated repression of PFKFB3. Arterioscler Thromb Vasc Biol. 2015;35(1):137–45.

42. Obach M, Navarro-Sabaté A, Caro J, Kong X, Duran J, Gómez M, et al. 6-Phosphofructo-2-kinase (pfkfb3) gene promoter contains hypoxia-inducible factor-1 binding sites necessary for transactivation in response to hypoxia. J Biol Chem. 2004;279(51):53562–70.

43. Min J, Zeng T, Roux M, Lazar D, Chen L, Tudzarova S. The Role of HIF1α-PFKFB3 Pathway in Diabetic Retinopathy. J Clin Endocrinol Metab. 2021;106(9):2505–19.

44. Reichard A, Asosingh K. The role of mitochondria in angiogenesis. Mol Biol Rep. 2019;46(1):1393–400.

45. Brix B, Mesters JR, Pellerin L, Jöhren O. Endothelial cell-derived nitric oxide enhances aerobic glycolysis in astrocytes via HIF-1α-mediated target gene activation. J Neurosci. 2012;32(28):9727–35.

46. Chen H, Fan W, He H, Huang F. PGC-1: a key regulator in bone homeostasis. J Bone Miner Metab. 2022;40(1):1–8.

47. Costa AD, Pierre SV, Cohen MV, Downey JM, Garlid KD. cGMP signalling in pre- and post-conditioning: the role of mitochondria. Cardiovasc Res. 2008;77(2):344–52.

48. Campbell JE, Drucker DJ. Pharmacology, physiology, and mechanisms of incretin hormone action. Cell Metab. 2013;17(6):819–37.

49. Baggio LL, Drucker DJ. Biology of incretins: GLP-1 and GIP. Gastroenterology. 2007;132(6):2131–57.

50. Vita JA, Treasure CB, Nabel EG, McLenachan JM, Fish RD, Yeung AC, et al. Coronary vasomotor response to acetylcholine relates to risk factors for coronary artery disease. Circulation. 1990;81(2):491–7.

51. Lee J, Bae EH, Ma SK, Kim SW. Altered Nitric Oxide System in Cardiovascular and Renal Diseases. Chonnam Med J. 2016;52(2):81–90.

52. Nisoli E, Falcone S, Tonello C, Cozzi V, Palomba L, Fiorani M, et al. Mitochondrial biogenesis by NO yields functionally active mitochondria in mammals. Proc Natl Acad Sci U S A. 2004;101(47):16507–12.

53. De Palma C, Falcone S, Pisoni S, Cipolat S, Panzeri C, Pambianco S, et al. Nitric oxide inhibition of Drp1-mediated mitochondrial fission is critical for myogenic differentiation. Cell Death Differ. 2010;17(11):1684–96.

54. Ma T, Du X, Pick JE, Sui G, Brownlee M, Klann E. Glucagon-like peptide-1 cleavage product GLP-1(9-36) amide rescues synaptic plasticity and memory deficits in Alzheimer’s disease model mice. J Neurosci. 2012;32(40):13701–8.

55. Tomas E, Stanojevic V, Habener JF. GLP-1-derived nonapeptide GLP-1(28-36)amide targets to mitochondria and suppresses glucose production and oxidative stress in isolated mouse hepatocytes. Regulatory peptides. 2011;167(2-3):177–84.

56. Bierhansl L, Conradi LC, Treps L, Dewerchin M, Carmeliet P. Central Role of Metabolism in Endothelial Cell Function and Vascular Disease. Physiology (Bethesda). 2017;32(2):126–40.

57. Eelen G, de Zeeuw P, Simons M, Carmeliet P. Endothelial cell metabolism in normal and diseased vasculature. Circ Res. 2015;116(7):1231–44.

58. Davidson SM, Duchen MR. Endothelial mitochondria: contributing to vascular function and disease. Circ Res. 2007;100(8):1128–41.

59. Limbourg A, Korff T, Napp LC, Schaper W, Drexler H, Limbourg FP. Evaluation of postnatal arteriogenesis and angiogenesis in a mouse model of hind-limb ischemia. Nat Protoc. 2009;4(12):1737–46.

